# Social isolation produces a brain-region specific expansion of microglia structure and reorganization of neural activity

**DOI:** 10.1101/2022.06.20.496842

**Authors:** Alex P. Vu, David Lam, Cayla Denney, Kelly V. Lee, Jason R. Plemel, Jesse Jackson

## Abstract

Social isolation is a profound form of psychological stress that impacts the mental health of a large proportion of society. Other experimental models of stress and injury have demonstrated microglia activation and alterations in neural activity. Microglia and neural activity undergo coordinated changes under physiological and pathological states. However, the effect of social isolation on microglia and neural activity has not been thoroughly investigated. Here we show that the dorsal medial hypothalamus and hippocampal CA2 region of male mice undergo an increased microglia volume and branching following social isolation, whereas females exhibit this increase in the hypothalamus only. The prefrontal cortex, central amygdala, nucleus accumbens shell, and visual cortex did not exhibit changes in microglia structure in either male or female mice. The home cage resting level of neural activity, as measured by the immediate early gene c-fos, was reduced in CA2 and the prefrontal cortex of female but not male mice following isolation. However, the co-variation in neural activity across brain regions was abolished in male but not female isolated mice. These data show that different brain regions undergo independent and dissociable changes in microglia structure and network activity following social isolation which may account for changes in cognition and behavior associated with this form of psychological stress.

## Introduction

Chronic social isolation is a pervasive problem in society, yet there is an incomplete understanding of how it affects the brain. Social isolation is psychologically stressful and is a causal and exacerbating factor for neuropsychiatric and neuroinflammatory conditions (Banerjee and Rai, 2020; Ieraci et al., 2016; Karelina et al., 2009; Kohn and Clausen, 1955; Taylor et al., 2018). Recent studies have shown that social isolation generates a host of behavioral deficits such as aggression, enhanced anxiety, and persistent fear responses, and these behavioral effects can be mediated by dissociable circuits throughout the brain (An et al., 2017; Hilakivi et al., 1989; Park et al., 2021; Weiss et al., 2004; Zelikowsky et al., 2018). Brain areas such as the central amygdala (Zelikowsky et al., 2018), prefrontal cortex, hippocampus (Hitti and Siegelbaum, 2014; Smith et al., 2016), and nucleus accumbens (Deutschmann et al., 2022; Park et al., 2021; Wallace et al., 2009) have a demonstrated importance in social behaviors, and these regions undergo cellular and neural activity changes following isolation (Cacioppo et al., 2015; Deng et al., 2019; Filipović et al., 2018; Stanisavljević et al., 2019; Zelikowsky et al., 2018).

Microglia are the resident immune cells of the brain, clearing cellular debris and pruning synapses as part of normal development (Paolicelli et al., 2011; Schafer et al., 2012; Sipe et al., 2016) and in response to injury and pain (Herzog et al., 2019; Neumann et al., 2009; Taylor et al., 2017). During disease conditions, microglia become reactive as defined by a biochemical or morphological divergence from a homeostatic, or surveillant, microglial state. While microglia reactivity is critical to protect brain tissue against injury or disease, under certain disease conditions reactive microglia can be detrimental (Block et al., 2007; Xu et al., 2016). In addition to seeing microglia reactivity during injury and disease, microglia can become reactive during chronic psychological stress (Hellwig et al., 2016; Tynan et al., 2010). For example, early life social isolation, where animals are isolated immediately following weaning, causes increased microglia reactivity initially with subsequent apoptosis, which is thought to be related to neuroinflammatory changes and oxidative stress (Gong et al., 2018; Jiang et al., 2013; Schiavone et al., 2009; Wang et al., 2017). However, the effect of adult social isolation on microglia and neural activity is less well documented.

Microglia are also responsive to neuronal activity changes and undergo enhanced branching and even become reactive in response to both neuronal hypoactivation and hyperactivation (Liu et al., 2019; Stowell et al., 2019; Umpierre et al., 2020). Microglia not only respond to neuronal activity, but they can actively regulate it. For example, microglia negatively regulate neuronal activity to prevent seizures (Badimon et al., 2020). As neural activity is responsible for signalling between brain regions, we were interested if social isolation alters both microglia structure and neural activity. We measured the changes in microglia branching and neural activity in response to social isolation in adult male and female mice. We focus on brain regions that have been previously shown to be affected by social isolation: medial prefrontal cortex (mPFC), shell of the nucleus accumbens (NAcc), CA2 of the hippocampus, central amygdala (CeA) and the dorsomedial hypothalamus (DMH) (Park et al., 2021; Stanisavljević et al., 2019; Zelikowsky et al., 2019). We found that the DMH and CA2 undergo increased microglia volume and branching. Neural activity was reduced in the CA2 and mPFC in female mice, whereas male mice had reduced neural activity correlations between brain regions (across mice). This work helps to further our knowledge of how different cell types and brain regions undergo alterations in structure and activity following social isolation.

## Methods and Materials

### Animal protocol

All procedures were performed according to the Canadian Council on Animal Care Guidelines and were approved by the University of Alberta Animal Care and Use Committee (AUP2711). Female (n=19) and male (n=24) adult (P60-120) C57BL/6 mice were obtained from Charles River Canada. Group housed mice were caged in sex-specific groups of 3-5 while isolation housed mice were separated into single housing cages for 14 days. Mice were housed in a temperature and humidity-controlled animal facility on 12h dark/light cycle with food and water ad libitum. Group housed and socially isolated mice were housed on the same rack in the animal facility.

### Perfusion and Tissue Sectioning

Mice were deeply anesthetized and transcardially perfused with ice cold phosphate buffered saline (PBS), followed by 4% paraformaldehyde (PFA) in PBS. Brains were extracted and postfixed in 4% PFA for 24–48 hours and stored in PBS at 4°C until sectioning. Brains were mounted in 2% agarose and sectioned at 50 μm using a vibratome (Leica VT1000s). Coronal sections were used for all brains. Slices were stored in antifreeze (50% PBS, 25% ethylene glycol, and 25% glycerol) at −20°C until immunohistochemistry was performed.

### Immunohistochemistry

Nine coronal slices from each brain were collected at anterior – posterior levels relative to bregma spanning in millimeters: +2, +1, +0.5, 0, −1, −1.5, −2, −2.5, −3. Antifreeze was washed off with 1X PBS (3 X 10 min) on a shaker, then incubated in blocking solution (2% Bovine Serum Albumin – 0.4% Triton X-100 in 1X PBS) on a shaker for 2 hours at room temperature. Slices were then incubated overnight in 4°C on a shaker in primary antibody solution: rabbit anti-cFos (1:1000), goat anti-Iba1 (1:500, Novus cat. # NB100-1028, RRID: AB_521594), in blocking solution. Slices were rinsed in 0.1% PBS-T (3 X 10 min) on a shaker then incubated for 4 hours at 4°C on a shaker in the dark, in secondary antibody solution: donkey anti-Goat Dylight 488 (1:500, Invitrogen), Donkey anti-rabbit Alexa 647 (1:500, Invitrogen), in blocking solution. Slices were then rinsed with 0.1% PBS-T (3 X 10min), then 1X PBS (2 X 10min) on shaker. They were then mounted on Fisher Superfrost Plus Microscope Slides and cover slipped with ProLong Gold antifade reagent with DAPI.

### Imaging

Six brain regions were identified for imaging: medial prefrontal cortex (mPFC), nucleus accumbens (NAcc), central amygdala (CeA), CA2 of the hippocampus (CA2), dorsomedial hypothalamus (DMH), and primary visual cortex (V1). Slides labeled with iba1 for microglia morphology were imaged on a Leica SP5 confocal microscope (DM4000) with a 40X objective (NA:0.8) with 1.5x zoom. Images were taken at a resolution of 1024×1024 pixels, and a 30μm Z-stack at 1μm z-step size. For neuronal activity quantification, c-fos-labeled slides were imaged on the Leica SP5 confocal microscope with a 20x objective. Images were taken at a resolution of 512×512 pixels. These images were taken using a 20μm Z-stack with a 5μm z-step size.

### Analysis

Experimenters were blinded to the identity of the animal during cell microglia analysis. All images were preprocessed in FIJI (Schindelin et al., 2012): scaled to 1559×1559 (to match 3D Morph’s preference of 0.166um/px), rolling background subtraction, noise de-speckled, and were thresholded to remove noise. Microglia structure within all images were then analyzed using 3D Morph (York et al., 2018) where the main outputs for microglia morphology were cell territory volume, cell volume, and endpoints. Between 10-25 cells were identified in each image of the same section and their values were averaged to represent a single animal.

Experimenters were blinded to the identity of the animal during cell counting. For c-fos quantification, images were preprocessed in FIJI (rolling background subtraction and brightness thresholding) and then had the number of c-fos positive cells counted in an image. To represent the brain region of one animal, the average of 2-3 images were used. We also analyzed the correlation coefficient of c-fos expression between regions across all animals within a given group housed or social isolated group, in order to determine the correlation in c-fos expression between two brain regions across mice (Wheeler et al., 2013). As we quantified c-fos in six regions, there were a total of fifteen pairwise correlation coefficient values that were visualized as a correlation coefficient matrix.

### Statistical Analysis

For statistical analysis, the mean and standard error of the mean were calculated within each brain region for the four groups (Male: group housed, Male social isolated, Female group housed, and Female socially isolated). A two-way analysis of variance using sex and isolation status as factors was conducted. Post-hoc follow up t-tests were conducted using the Sidak’s correction for multiple comparisons. For t-tests unpaired t-tests that did not assume equal variances between groups were used to compare the means of social-housed versus isolated mice. All analyses were done in GraphPad Prism 8 (GraphPad Software, San Diego, California USA).

## Results

Although microglia have been examined in the context of psychological stressors in the past (Hellwig et al., 2016; Kreisel et al., 2014) it is unclear if there are brain region and sex-dependent changes in the morphology of microglia in response to adult social isolation. Iba1 was used to identify microglia within the brain tissue (Ito et al., 1998). We performed high resolution reconstruction of microglia in different brain regions previously shown to play a role in social behaviors (mPFC, NAcc, CeA, CA2 and DMH; (Stanisavljević et al., 2019; Zelikowsky et al., 2019) (**Figure 1A-C**). Primary visual cortex (V1) was also examined since we predicted there would be less change in this region as it is not known to participate in social behavior. Using open-source software (3D-morph), we measured the spatial territory, the volume, and the number of end points for each cell. In total we reconstructed 4170 microglia across all brain regions (**Figure 1B**) from a total of 43 mice (14 male group housed, 10 male socially isolated, 10 female group housed, and 9 female socially isolated).

**Figure 1:**
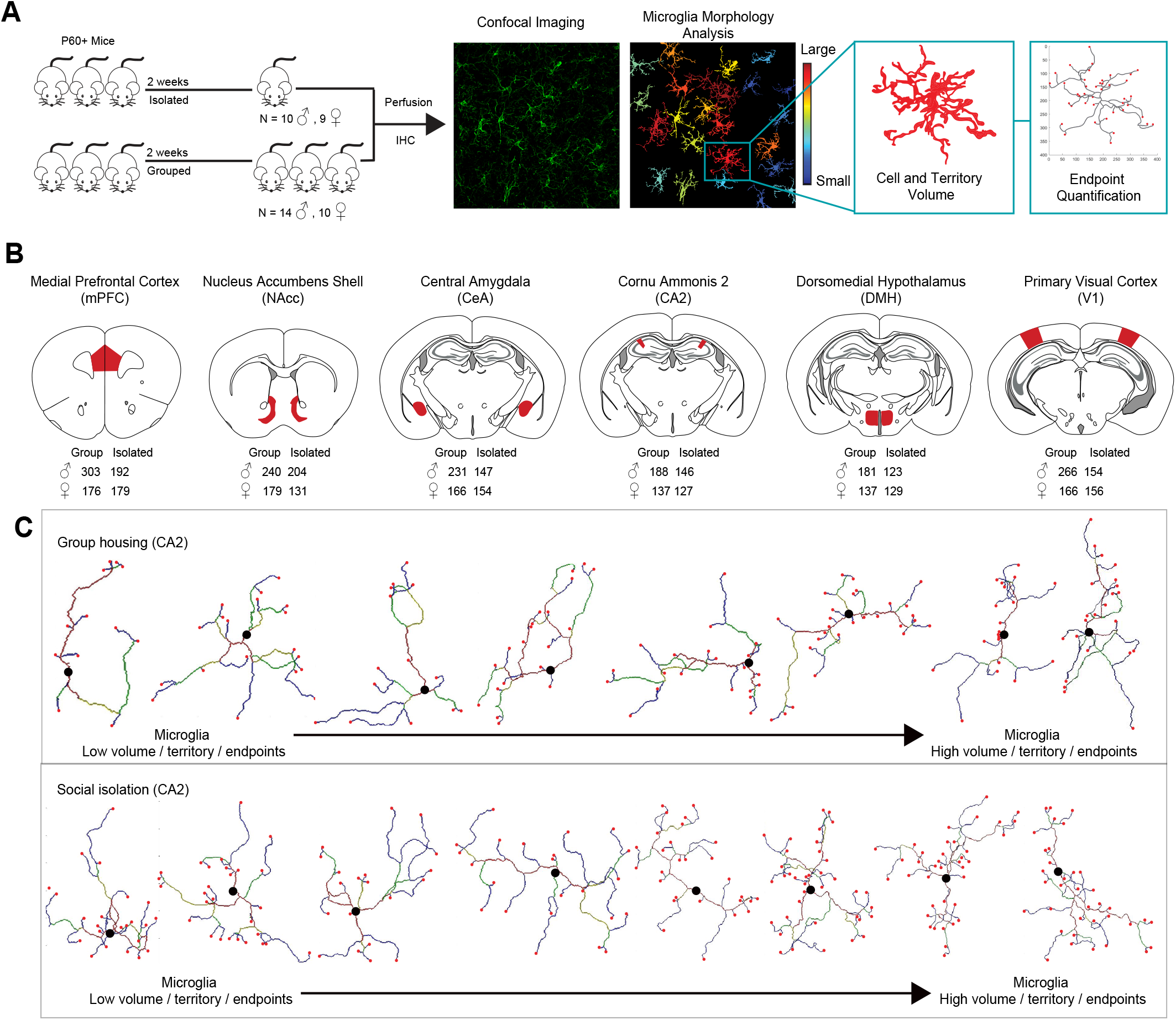
Experimental setup, brain regions analysed, and example data. **A**: Adult male and female mice were socially housed or isolated for two weeks, followed by perfusion and immunohistochemistry. An example field of view is shown of a maximum intensity projection image from Iba1 immunohistochemistry, followed by morphology analysis (color coded according to cell volume). The skeletonized microglia are binarized and spatially dilated for visualization. On the right is an expanded cell, showing the cell volume, and endpoints (red points). **B**: Brain atlas schematics showing the regions analyzed, highlighted in red. The total number of microglia in each group is indicated below. **C**: Example microglia reconstructions from the CA2 region of a male group housed mouse (top), and socially isolated male (bottom), arranged to highlight cells with low (left) to high (right) cell volume, spatial territory, and endpoint number.

Following two weeks of social isolation, we found that microglia in the DMH had significantly increased microglia territory, cell volume and endpoints per microglia within both male and females relative to group housed mice (**Figure 2A**, **Table 1**). Additionally, we saw this same change in microglia morphology male mice in CA2, but not in females (**Figure 2B**, **Table 1**). There were no significant differences in microglia morphology within the mPFC, CeA, NAcc or V1 between socially isolated and group housed mice in either male or female mice (**Figure 2C-F**). The CA2 and DMH exhibited a significant effect of sex on microglia structure, but there was no significant interaction between sex and isolation status (group versus single housed) (**Table 1**).

**Figure 2:**
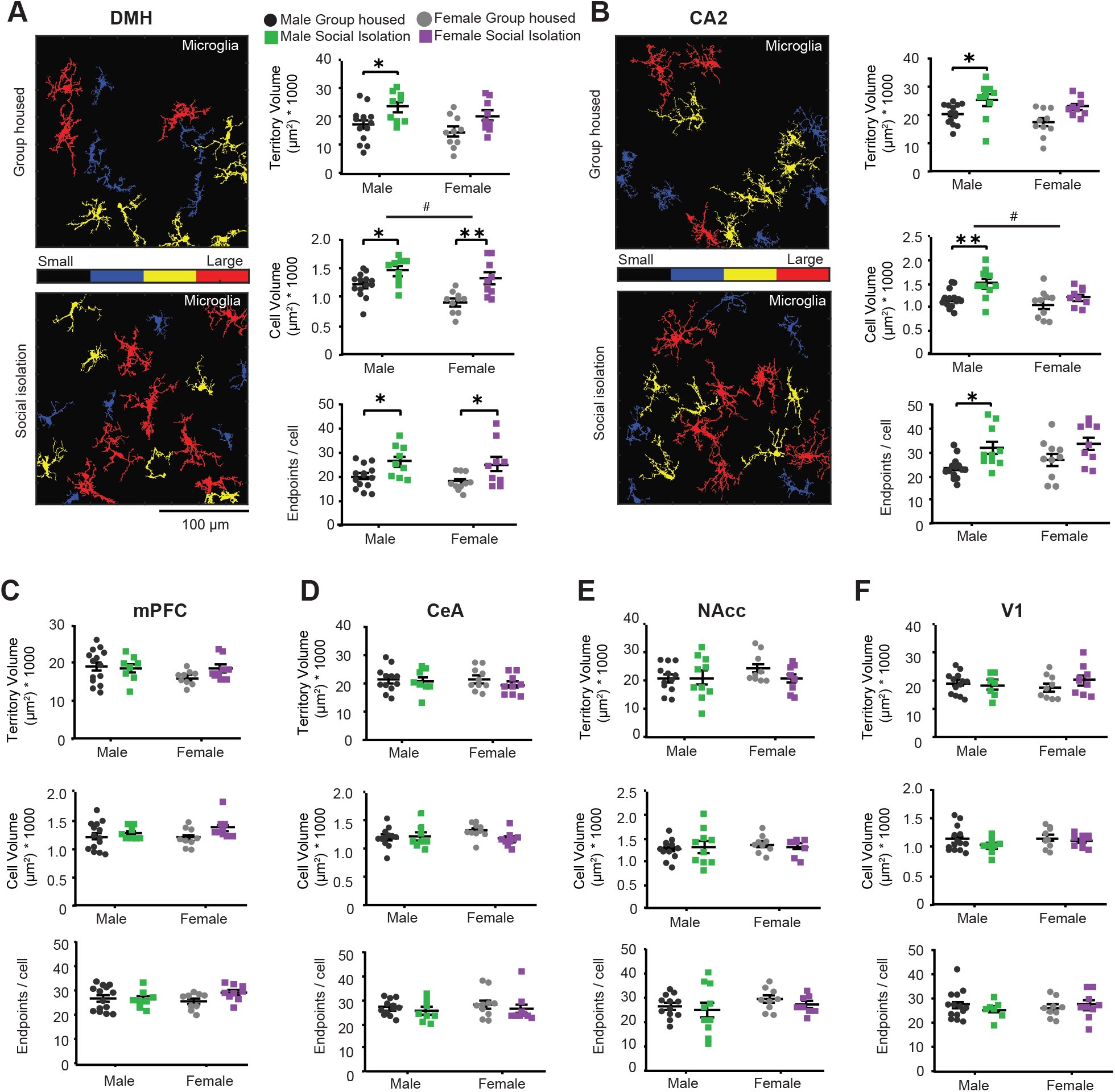
Region specific increases in microglia morphology following adult social isolation. **A**: Example zoomed in fields of view showing microglia in the DMH in group housed (top) and socially isolated female mice (bottom). The quantification of cell volume, territory, and endpoints/cell are shown on the right. Group identification is listed above and labelling strategy applies to all subpanels in the figure. Individual points indicate the mean of each mouse, and the error bars show the standard error of the mean. **B – F**: the same as A but for CA2, mPFC, CeA, NAcc, and V1 respectively. * p <0.05, ** p <0.01. See Table 1 for a complete list of descriptive statistics and comparisons.

**Table 1:**
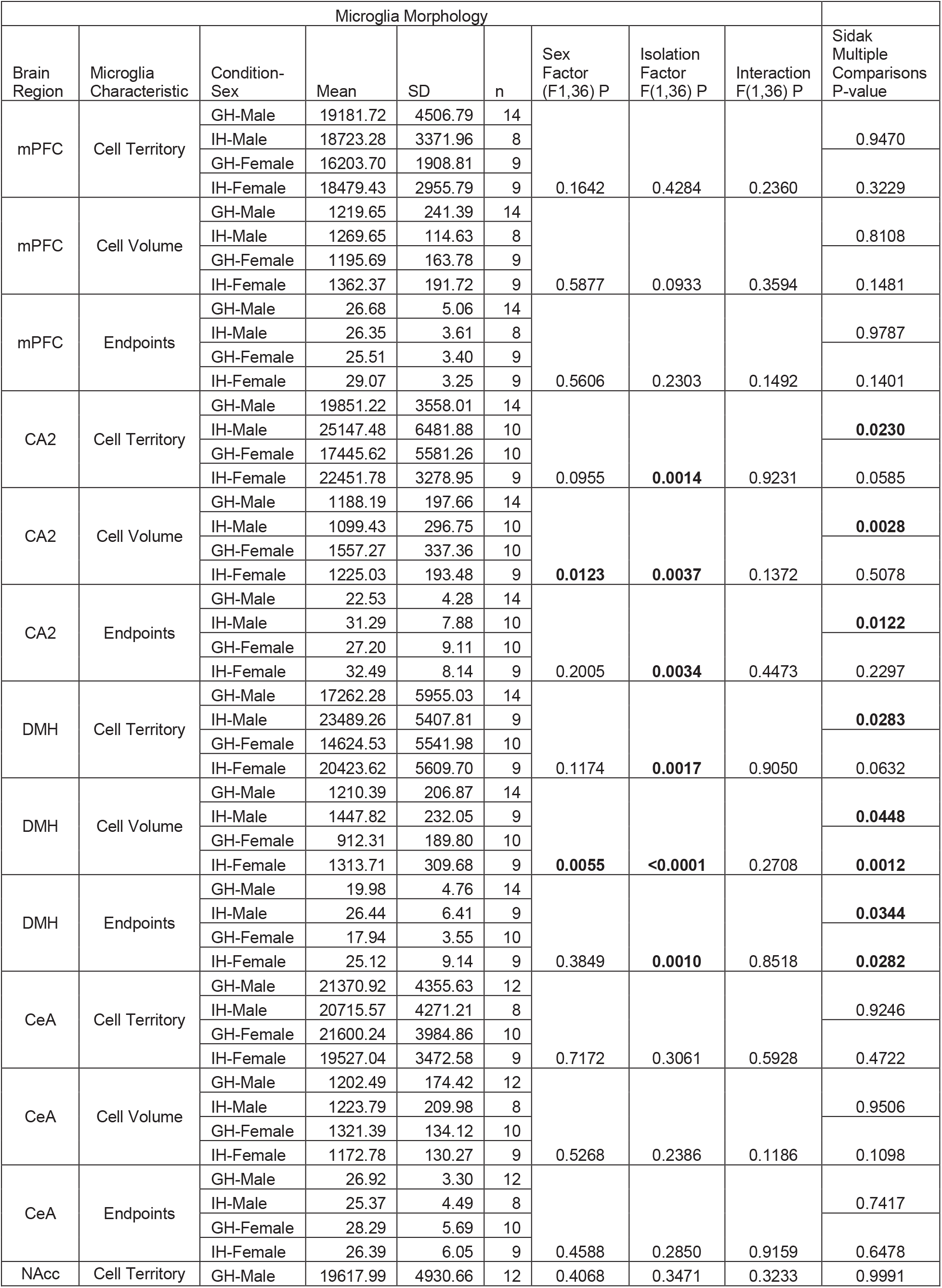
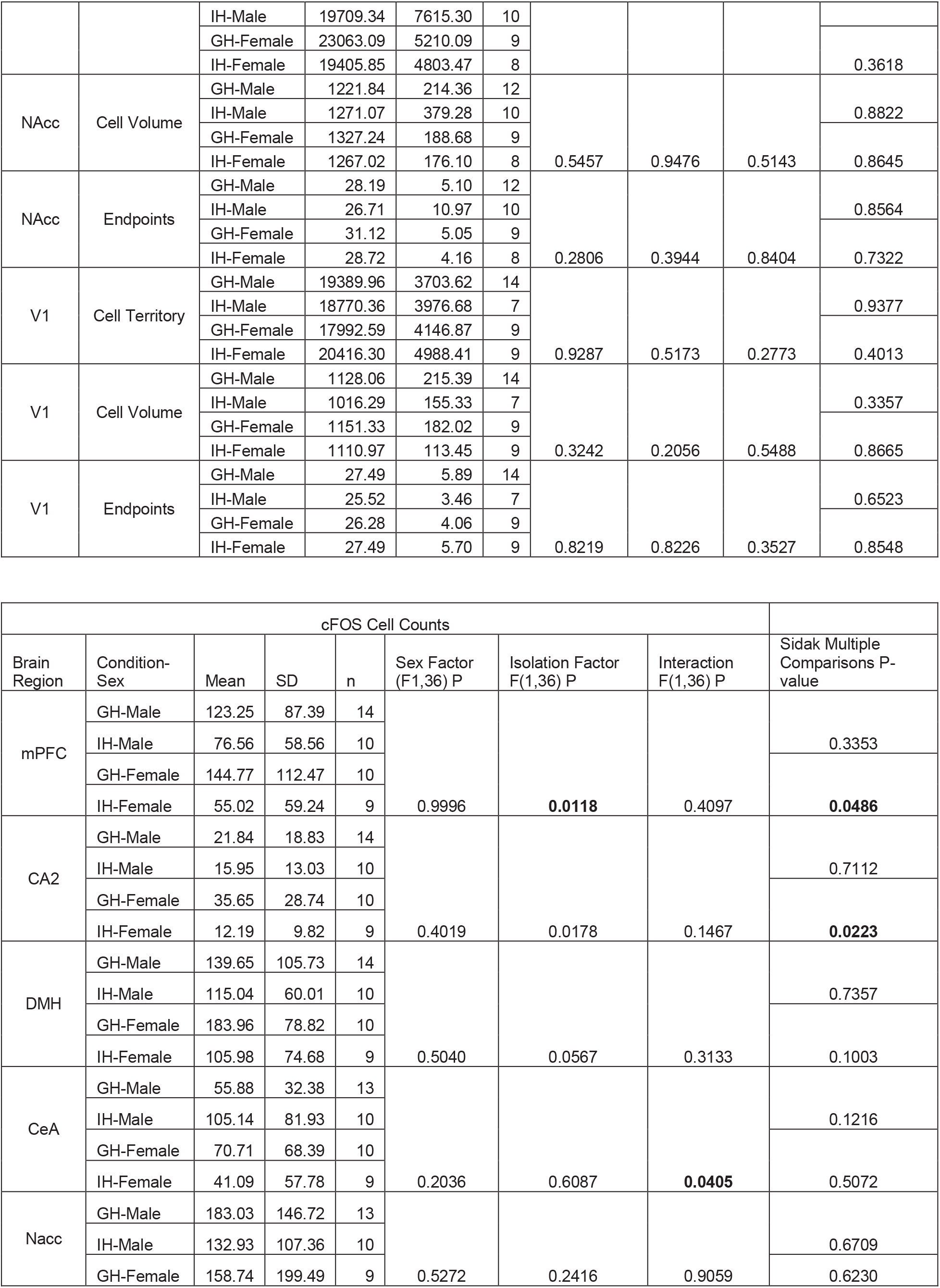
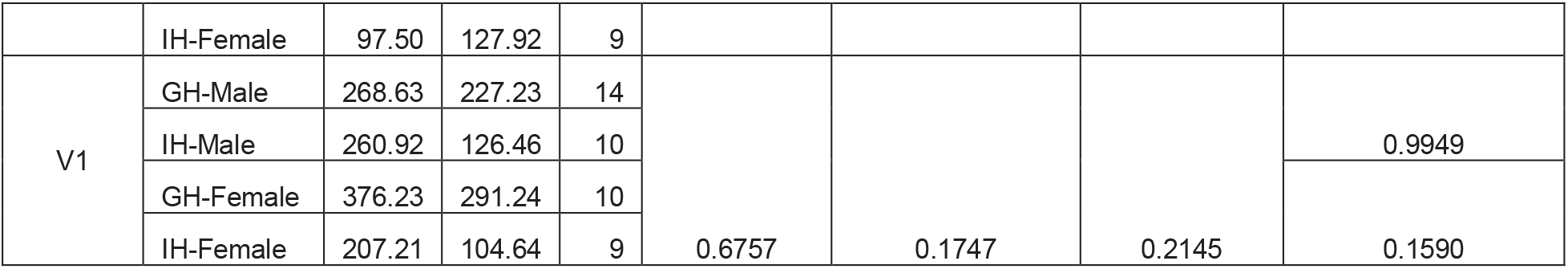
Descriptive statistics and pairwise comparisons from all data shown in Figure 2 and Figure 3.

We next wanted to determine if isolated mice also showed differences in basal neuronal activity. To do this we used c-fos immunohistochemistry as a marker for neural activity. Previous research has shown that c-fos expression changes during social isolation in some cortical and subcortical brain regions (Stanisavljević et al., 2019; Zelikowsky et al., 2019; Zorzo et al., 2019). However, we wanted to determine if c-fos changes occurred in the same brain region where we see observe differences in microglia morphology (**Figure 2**). Only the mPFC and CA2 showed a small decrease in c-fos in female isolated mice but this effect was not observed in males (**Figure 3A-B**). Finally, we measured how c-fos activity covaried between brain regions in group housed and socially isolated mice. For this analysis the Pearson correlation coefficient was calculated between c-fos measurements for each pair of brain regions, across mice. The brain regions analyzed are part of a group of stress and socially responsive brain regions (aside from V1), and therefore c-fos levels were suspected to co-vary between brain regions within a given mouse. As expected, group housed males had a c-fos activity correlation between regions of 0.46±0.19 (across 15 pairs of brain regions) which was significantly reduced to −0.08±0.25 with social isolation (p = 1.8×10^−5^), suggesting a lower functional connectivity in males (**Figure 3D**). However, female mice did not show a change in inter-region c-fos correlation (r = 0.47±0.27 versus 0.32±0.28, p = 0.15) (**Figure 3E**).

**Figure 3:**
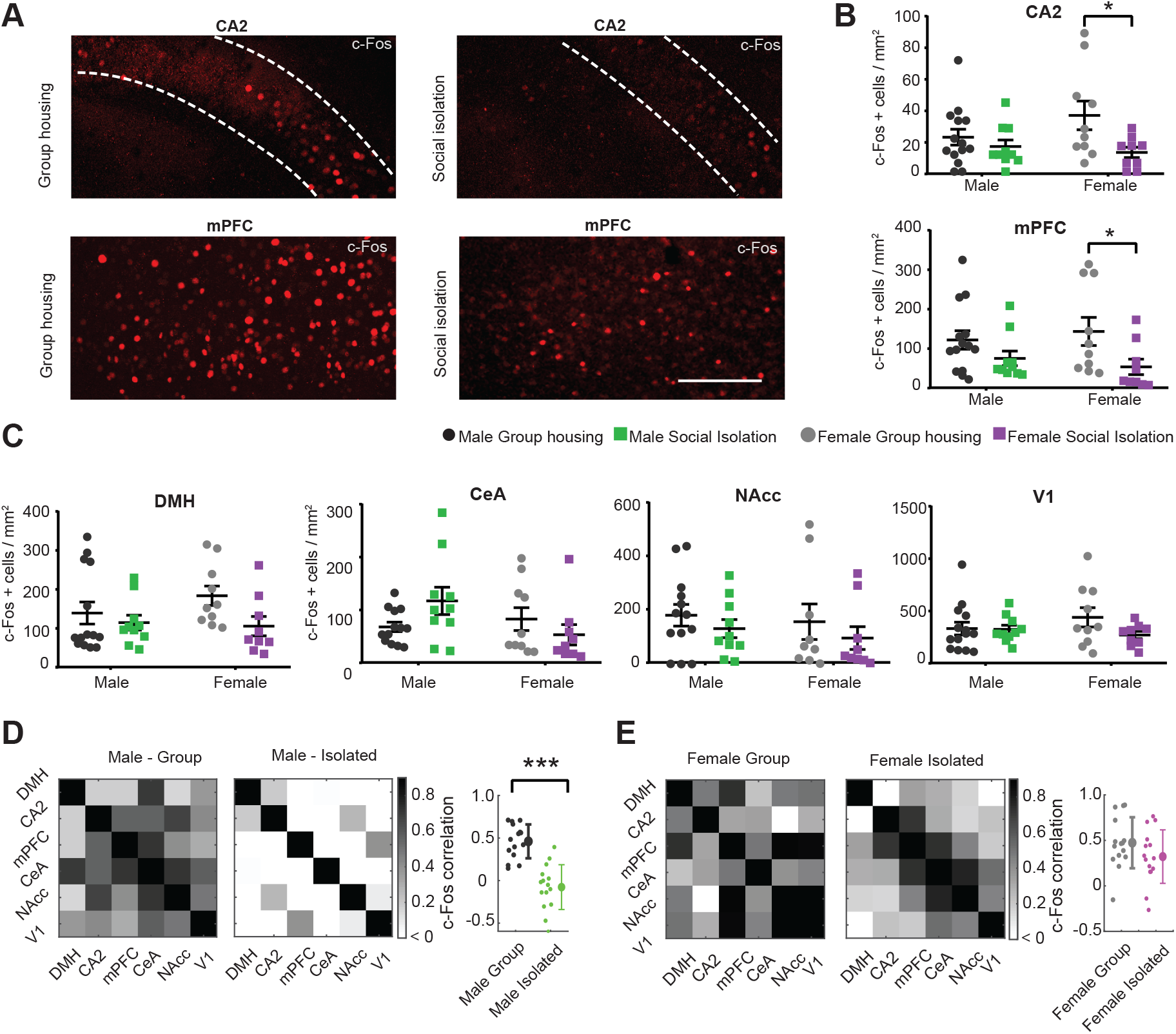
Social isolation produces a reduction of c-fos activity in the female mice and inter-region activity correlations in male mice. **A**: Example images showing c-fos the CA2, and mPFC of female group housed and socially isolated mice. **B**: Quantification of c-fos density in the CA2 and mPFC in each brain region in all groups. **C**: c-fos density in other brain region was not statistically altered. **D**: Left: correlation matrices plotting the Pearson correlation coefficient of c-fos activity levels between brain regions in group house and isolated mice. Matrices are symmetrical across the diagonal. Right: The interregional c-fos correlations are also shown as a scatter with the mean ± standard deviation. **E**: The same as **D**, but for females. Scale bar in A is 0.1mm, * p <0.05, *** p <0.001. See Table for a complete list of statistics.

## Discussion

In the present study we have shown that social isolation stress results in significant increases in microglial branching and territory occupancy in brain regions associated with social behaviors such as the DMH and CA2. Decreases in c-fos were also detected in females in the mPFC and CA2 brain regions, but not in males. However, correlated activity between brain regions was reduced in males and preserved in females. These data compliment previous studies showing microglia undergo morphological changes under various forms of stress including post weaning social isolation, physiological restraint, or chronic social defeat (Gong et al., 2018; Lehmann et al., 2019; Tynan et al., 2010).

One potential explanation underlying the morphological expansion of microglia seen in chronic social isolation is that psychological stress induces an inflammatory response within the brain, likely driven by microglia (Calcia et al., 2016; Hellwig et al., 2016; McKim et al., 2016; Niu et al., 2020; Piirainen et al., 2017). Such an immune response, due to chronic stress, may have parallels to inflammatory mechanisms related to psychiatric disorders such as depression, anxiety and schizophrenia (Müller, 2018; Rooney et al., 2020; Torres-Platas et al., 2014; Wang et al., 2018), all of which have also been linked to social isolation (Ieraci et al., 2016; Kohn and Clausen, 1955; Lee et al., 2021; Taylor et al., 2018). However, a related possibility is that neurons within socially related brain regions become hypoactive following social isolation, and this hypoactivity produces an increase in microglia process extension. Consistent with this possibility, experimentally-induced suppression of neuronal activity in the neocortex results in enhanced microglial branch extension (Liu et al., 2019; Stowell et al., 2019; Umpierre et al., 2020). It is known that microglia and neurons communicate bidirectionally, and therefore it remains to be determined if microglia cause changes in neural activity, if neural activity causes the changes in microglia, or if these under these conditions these two phenomena are not causally linked.

Social behavior is controlled by a spatially distributed set of brain regions. We analyzed five brain regions known to participate in social behaviors (mPFC, NAcc, CA2, CeA and DMH)(Park et al., 2021; Stanisavljević et al., 2019; Zelikowsky et al., 2019), together with visual cortex (no known role in social behavior). Recent work demonstrated that maladaptive cellular changes in separate socially-related neural circuits are responsible for different behavioral deficits arising following social isolation (Matthews and Tye, 2019; Zelikowsky et al., 2018). For example, increased aggression is attributed to changes in the hypothalamus (Brasil et al., 2020; Cui et al., 2013; Zelikowsky et al., 2018), whereas social memory may arise from changes in the hippocampus (Deng et al., 2019; Kogan et al., 2000; Smith et al., 2016). We also observed changes in the DMH and CA2, suggesting that perhaps microglia changes may in some way contribute to both the increased aggression and decreased social memory deficits in socially isolated mice. The mPFC and NAcc also have important roles in the encoding and recognizing social stimuli (Murugan et al., 2017). However, we did not observe changes in microglia structure or basal activity rates in these regions, suggesting that any deficits in function within these circuits are independent of microglia specific changes (Park et al., 2021). We did not detect a change in c-fos activity in any brain region within males, whereas several prior studies have shown reductions in c-fos in the mPFC, NAcc, and hippocampus. One possibility for this discrepancy is that we analyzed c-fos at ‘rest’ in the home cage, whereas prior studies measured c-fos following behavior.

In addition to finding region-specific changes in microglia morphology we also found some subtle differences between males and female mice. In male mice, microglia displayed more pronounced changes in CA2 whereas the DMH showed changes in both males and females. On the other hand, females showed a decrease in neural activity, whereas males did not. However, the c-fos correlation between brain regions was reduced in male mice but not females, suggesting that perhaps communication between brain regions is more affected in males than females. Supporting this data, another recent report also demonstrated a breakdown of functional connectivity in mice using wide-field cortical imaging (Kim et al., 2021), but the sex of the mice was not reported. Sex dependent (and divergent) changes have been shown in anxiety and anhedonia behaviors following social isolation (Oliver et al., 2020; Ross et al., 2019; Weiss et al., 2004) and in psychiatric diseases such as depression (Bangasser and Cuarenta, 2021; Kessler et al., 2012; Labaka et al., 2018). Given the link between social isolation, microglia and the formation of these diseases, further research is required to explore how social isolation affects the brain in a sex dependent manner and what role microglia plays in this brain reorganization.

It is now well understood that microglia play a considerable role responding to and regulating neural activity under physiological and pathological states such as stress. Social isolation is a strong psychological stressor and risk factor for a wide range of other diseases. The results of the present study show sex- and region-specific microglia changes in morphology and neuronal activity in the context of adult social isolation. Given these findings, further study is required to determine if and how the dynamic interplay between microglia and neural activity contribute to different forms of cognitive and behavioral impairment in social isolated animals.

## Acknowledgements

We thank The Faculty of Medicine & Dentistry Cell Imaging Center including Steve Ogg, and Greg Plummer for assistance with microscopy, and Dr. Anna Taylor for comments and advice on the manuscript. Funding was provided by Canada Foundation for Innovation John R. Evans Leaders Fund (JELF), Grant/Award Number: 37931; Canadian Institutes of Health Research, Grant/Award Number: 426485; National Alliance for Research on Schizophrenia and Depression; Natural Sciences and Engineering Research Council of Canada, Grant/Award Number: RGPIN2018-05212.

